# Bacterial-tyramine-induced renal-gut countercurrent flow regulates gut immune response by promoting ROS accumulation and gut peristalsis in the fly *Bactrocera dorsalis*

**DOI:** 10.1101/2024.03.22.586092

**Authors:** Yanning Liu, Rengang Luo, Bruno Lemaitre, Hongyu Zhang, Xiaoxue Li

**Author notes:** **Author for correspondence:** Email: Xiaoxue Li (Lead contact) Email: Hongyu Zhang.

## Abstract

Insects increase gut peristalsis to purge ingested pathogens. They also rely on the Duox-ROS system to mount an efficient gut response. However, how these two host defense mechanisms are coordinated to clear pathogens remained unclear. We found that gut peristalsis is the major force for bacteria elimination, while Duox is necessary as a signaling molecule to activate gut peristalsis in a Tephritidae fruit fly *Bactrocera dorsalis*. We found an insect-conserved renal-gut countercurrent flow formed upon recognizing bacteria-derived tyramine counteracts the gut peristalsis forced defecation and drives the accumulation of ROS. Furthermore, we demonstrated that the renal-gut countercurrent flow maintains proper microbiota composition. Our work provided a novel renal-gut interaction that ensures an efficient gut-immune response.

## Introduction

Living in a microbe-rich environment, insects such as Dipteran interact with a broad range of symbionts, opportunists, and pathogens. This diversity of cohabitants has likely shaped the sophisticated insect immune system, that must be capable to combat pathogens while promoting the growth of beneficial microbes. Studies mostly done in *Drosophila* have shown the multiple roles of the gut microbiota on insect health by promoting host development under poor nutrient conditions, providing vitamins, or preventing infection^1–3^. Dysregulation of the gut microbiota affects gut homeostasis and shortens lifespan^1, 4^. These studies have also revealed that the gut immune response of insects is complex, encompassing various constitutive and inducible defenses.

First, physical barriers protect the gut epithelium from direct contact with pathogens. These include the cuticle that lines the foregut and hindgut, and a chitinous layer called the peritrophic membrane (PM) that lines the midgut^5, 6^. The PM forms a porous matrix permeable enough to allow the digestive process while separating bacteria and harmful particles from the gut epithelium^7^. An acidic region in the middle part of the midgut also contributes to eliminating ingested bacteria^8, 9^. Second, the mechanical flushing of pathogens out of the intestine by peristalsis is thought to be a vital host defense mechanism. Peristalsis involves a wave-like longitudinal and circular muscle contraction that moves food and bacteria through the digestive tract^10^. The waves can be short, regional reflexes or long, continuous contractions that travel the whole gut. In insects, as in mammals, gastrointestinal motility is controlled by enteroendocrine cells as well as enteric and central nervous systems ^11, 12^. In *Drosophila*, ingestion of bacteria promotes strong midgut visceral contractions. a process involving ROS production by the NADPH Duox by enterocytes^13^. ROS then binds to the TRPA1 receptor found in enteroendocrine cells, which secrete the diuretic hormone DH31. DH31 binds to its receptor in neighboring visceral muscles, triggering contractions^14^.

Production of antimicrobial peptides by the Imd pathway and ROS by the NADPH Duox provides two complementary immune inducible mechanisms. Bacteria ingestion triggers the expression of several antimicrobial peptide genes in specific domains along the digestive^15^. This response is regulated by the Imd-NF-ΚB pathway upon the sensing of peptidoglycans released by bacteria via receptors of the PGRP family^15–19^. Ingestion of bacteria also triggers the production of possibly microbicidal hypochlorous acid (HOCl) by Duox. Its microbidical role is thought to be crucial for *Drosophila*’s ability to survive oral bacteria infection^20^. Production of ROS by Duox is achieved through the recognition of pathogenic bacteria released by uracil, which stimulates the Gαq-PLCβ-Ca_2_^+^ pathway, increasing the enzymatic activity of Duox^20–22^. In addition, the sequential activation of MEKK1-MKK3-p38 MAPK and lipid catabolism also upregulated transcription of the *Duox* gene^23, 24^.

While initial studies have shown that Duox-derived HOCl has a direct bactericidal effect, other studies revealed another role for Duox in gut homeostasis. As described above, Duox is also involved in peristalsis but also contributes to the increased stem cell proliferation observed upon bacterial infection^25^. Moreover, we have recently shown that Duox in the Malpighian tubules is required for gut homeostasis post oral infection^26^. Malpighian tubules form the insect kidney, filtering the hemolymph and regulating osmolarity/water homeostasis^7^. Malpighian tubules are connected at the posterior end of the midgut, where they generate primary urine^27^. Studies done in several insect species, notably locusts, have revealed the existence of counterflows that can flush Malpighian tubule liquid toward the anterior parts of the gut^28, 29^. The existence of retrograde fluid flow in the insect gut is possibly due to the peritrophic matrix that transversally compartmentalizes the midgut. According to this model, food and bacteria transit forward along the gut in the endoperitrophic space formed by the PM, while the countercurrent flow initiated by Malpighian tubules takes place in the ectoperitrophic space between the PM and the epithelium. We have recently shown that this counterflow is also observed in *Drosophila melanogaster* ^26^. Our study reveals that this countercurrent flow increases upon bacterial oral infection and this counterflow brings the tubules-produced JAK-STAT pathway secreted ligand Upd3 to the anterior part of the midgut to promote an increased epithelium renewal. Thus, this study reveals an additional role of Duox in midgut homeostasis and indispensable role of the tubules in gut immunity. The observation that this counterflow is increased upon bacterial infection suggested that it could play an important role in host defense. Collectively, all these studies have highlighted diverse roles of Duox mediated production of ROS in the gut either as direct microbicidal agent or as indirect signaling molecules promoting peristalsis and countercurrent flow. It is however unclear how Duox mediate bacteria clearance upon gut infection.

In this study, we aim at clarifying the role of Duox-ROS system in insect gut defense. We used the Tephritidae fruit fly *Bactrocera dorsalis,* an invasive dipteran pest feeding on over 250 vegetables and fruits, causing enormous economic losses worldwide. *B. dorsalis* larva lives in rotten fruits and owns a complex gut microbiome, making it a good model for gut immunity study^30^. Here, we showed that infection increased gut peristalsis to flush bacteria, a process that involves ROS produced by Duox. Importantly, the level of ROS in the midgut is maintained by the existence of countercurrent promoting ROS recycle/accumulation. We also show that this countercurrent flow requires the aquaporin Prip and is triggered upon the recognition of microbial-derived tyramine. Collectively, our study reveals the critical role of Duox in intestinal immunity by promoting gut peristalsis rather than by a direct microbicidal effect. Our study also revealed the importance of MpT-gut countercurrent flow to flush pathogenic bacteria out and plays a crucial role in gut microbiota maintenance.

## Results

### Bacterial infection triggers a countercurrent flow in *B. dorsalis*

We have recently shown that oral infection activates a renal-gut countercurrent flow in *Drosophila* that plays an important role in gut immunity by mediating gut epithelial renewal ^26^. We set up an *ex vivo* assay to test the existence of countercurrent flow in *B. dorsalis*. Briefly, flies were subjected to either oral infection for two hours with a gram-negative bacteria, *Providencia rettgeri,* or a control sucrose solution, then switched to fresh fly food (Supplementary Figure 1). In this way, we could observe the outcome of a short bacterial infection rather than a continuous oral infection. The contact gut and MpT were dissected and placed in the adjacent wells soaked in saline buffer and Schneider’s insect medium. The well with MpT was filled with additive Brilliant blue FCF as an indicator. So, if there is a renal-gut countercurrent flow, the dye in the MpT well will be absorbed and transferred to the midgut lumen in the separating wall (Figure 1A). The results showed that *P. rettgeri* infection induced the rapid MpT absorption of Brilliant blue dye, consistent with the existence of a countercurrent flow 2 h post infection. We observed the dye moved forward from the MpT lumen to the midgut and accumulated within the whole gut, including the anterior midgut and posterior midgut, and that this effect was stronger after infection with *P. rettgeri* (Figure 1B). We noticed an increase in the percentage of the gut displaying dye from 33.3% to 65.6% upon oral infection. Notably, almost half of the gut has a strong dye signal in the anterior part, suggesting a more robust flow than the unchallenged condition (Figure 1C). Moreover, we also saw the countercurrent flow positive gut in the unchallenged condition, suggesting this flow might have a role even without infection (Figure 1C). To further confirm our finding, we injected the neuropeptide Leucokinin (Kinin), one major hormone regulating MpT stellate cells’ secretion ability and monitored countercurrent flow. Strikingly, we observed that injection of Leucokinin stimulate countercurrent flow to the same extent than *P. rettgeri* infection. We observed the accumulation of the blue dye in the gut, particularly in the anterior midgut, demonstrating MpT activate this gut countercurrent flow (Figures 1D and E). Altogether, these data revealed the existence of an MpT-gut countercurrent flow after oral infection in *B. dorsalis*.

**Figure 1.**
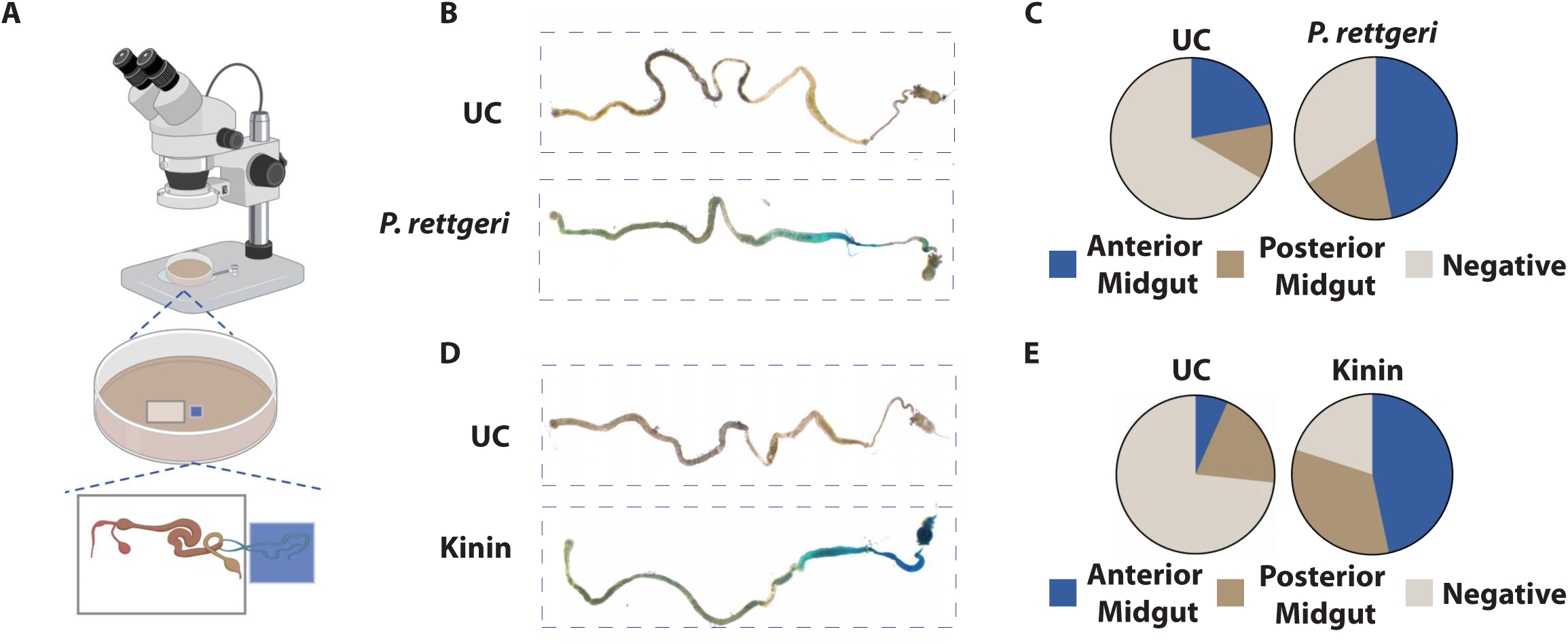
Bacterial infection triggers a countercurrent flow in *B. dorsalis*. **(A) Schematic diagram of *ex vivo* intestinal countercurrent detection method.** Two adjacent wells in the PDMS plate were filled with cell culture. The connected gut and MpT were put into two adjacent wells respectively, and the MpT well was supplemented with 0.05% bright blue solution. A drop of mineral oil was added to the junction of the two wells to cover the tubules preventing tissue dryout. **(B and C) *P. rettgeri* gut infection promotes gut countercurrent flow.** Oral infection induced a strong MpT-gut countercurrent flow indicated by the accumulation of blue dye in the anterior midgut (B) and quantification in (C). (n=36 for UC, n=32 for *P. rettgeri*). UC: unchallenged. **(D and E) Kinin promotes gut countercurrent flow generation.** Kinin injection induced an MpT-gut countercurrent flow similar to bacterial infection (D) and quantification in (E). (n=15 for imidazole, n=32 for kinin). UC: imidazole.

### Countercurrent flow in *B. dorsalis* requires the Aquaporin Prip

In *D. melanogaster*, aquaporin Drip and Prip in MpT stellate cells mediate rapid water flux^31^. We then explored the role of these aquaporins in controlling MpT-gut retro-flow in *B. dorsalis*. We confirmed that *Drip* and *Prip* genes were expressed in *B. dorsalis* MpT tubules (Supplementary Figures 2A and B). We found that *P. rettgeri* infection did not induce upregulation of these two genes (Supplementary Figures 2C and D). To study their function in countercurrent flow generation, we knocked down *Prip* and *Drip* using RNAi by injecting dsRNAs (Supplementary Figures 2E and F). *In vitro* countercurrent flow assay confirmed the role of *Prip* but not *Drip* in countercurrent flow formation. The results showed that countercurrent flow could reach the anterior midgut in *egfp* RNAi control flies but not in *Prip* RNAi flies after *P. rettgeri* infection, demonstrating its role in MpT-gut countercurrent flow formation (Figures 2A and B). However, we saw no difference in dye accumulation between the *Drip* RNAi and the control groups, suggesting *Drip* is not required for countercurrent flow formation (Figures 2C and D).

**Figure 2.**
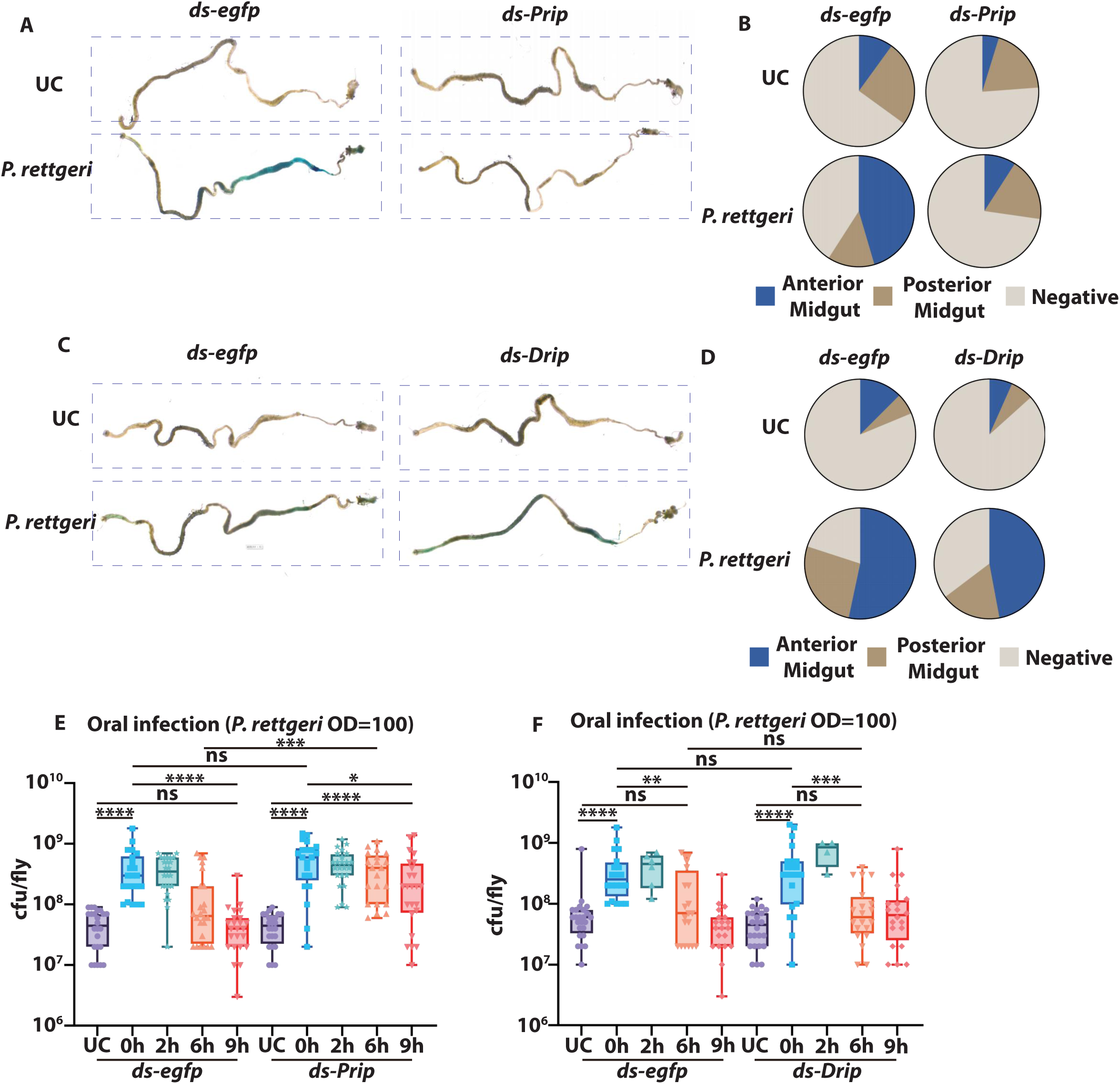
Prip-mediated MpT-gut countercurrent flow ensures bacteria elimination. **(A and B) *Prip* is required for gut countercurrent flow generation.** *Prip* RNAi flies showed very weak blue dye accumulation in the anterior midgut (A) and quantification in (B). (n=20-33). *egfp* RNAi flies were used as control. UC: unchallenged. **(C and D) *Drip* RNAi did not affect gut countercurrent flow generation.** There was no significant difference regarding blue dye accumulation in the gut between *Drip* RNAi and *egfp* RNAi flies indicated by countercurrent flow assay (C) and quantification in (D). *egfp* RNAi flies were used as control. The data represents the average of 3 biological replicates (n=15-17). **(E) *Prip* RNAi decreased the flies’ ability to eliminate gut invading bacteria.** *egfp* RNAi and *Prip* RNAi flies showed significant cfu counts differences at 6 h after the infection. *Prip* RNAi flies could not eliminate invading bacteria at 9 h after infection. *egfp* RNAi flies were used as control. The data represents the average of 3 biological replicates (n=21-24). UC: unchallenged. **(F) *Drip* RNAi did not affect bacteria elimination.** *Drip* RNAi did not affect the flies’ ability to eliminate gut-invading bacteria. *egfp* RNAi flies were used as control. The data represents the average of 3 biological replicates (n=4-24). UC: unchallenged. ** p<0.01, *** p<0.001, **** p<0.0001, ns non-significant, Mann Whitney test. UC: unchallenged.

### Countercurrent flow promotes bacterial clearance in *B. dorsalis*

The observation that counterflow is induced upon bacterial infection points to its role in host defense. Next, we investigated whether this MpT-gut retro flow is required for bacteria clearance. Both *Prip*, *Drip* RNAi flies and control *egfp* RNAi flies were fed with *P. rettgeri* for 2 hours and then switched to fresh food. We monitored bacterial persistence in the gut for the next 9 hours. Importantly, we found that *Prip* RNAi flies lost their ability to clear invading bacteria because these RNAi flies did not clear bacteria even after 9 h of infection (Figure 2E). However, *Drip* RNAi flies, which have normal retro-flow, have successfully cleared invading bacteria 6 h after *P. rettgeri* infection, showing no difference compared to the control group (Figure 2F). As Prip but not Drip is involved in the countercurrent flow formation, our results points to an important role of MpT-gut retro-flow in bacterial clearance. In conclusion, the above results demonstrated that the aquaporin *Prip*-mediated MpT-gut countercurrent flow is required for efficient bacteria clearance after oral infection in *B. dorsalis*.

### Exogenous tyramine is a signaling molecule that triggers MpT-gut countercurrent flow

The biogenic amine tyramine regulates many aspects of animal physiology. Tyramine also controls renal function in *Drosophila*^32^. We hypothesized that tyramine might be the signaling molecule inducing the countercurrent flow from the MpT to the gut. To test this hypothesis, we first knocked down *TyrR* in *B. dorsalis*. TyrR is the tyramine receptor regulating MpT stellate cell activity^33^. We monitored MpT-gut countercurrent flow using *in vitro* countercurrent experiments. *TyrR* knockdown strongly decreased the intensity of this retro-flow, as the dye could not reach the anterior midgut in most *TyrR* RNAi flies after infection (Figures 3A and B). More importantly, *TyrR* RNAi flies also had a weaker ability to clear invading bacteria (Figure 3C). These data indicated that tyramine is the signaling molecule that induces the MpT-gut countercurrent flow. To further strengthen our conclusions, we directly fed *B. dorsalis* with tyramine and detected countercurrent flow. The results showed that feeding flies with a diet supplemented with tyramine enhanced gut countercurrent flow, promoting dye accumulation in the anterior midgut, similar to the oral infection (Figures 3D and E). We also speculated whether feeding a diet supplemented with tyramine in *B. dorsalis* would enhance the ability to remove pathogenic bacteria. Feeding tyramine significantly increased flies’ ability to eliminate invading bacteria. Tyramine-fed flies eliminated invading bacteria within 1 h after infection compared with 2 h in the control flies (Figure 3F).

**Figure 3.**
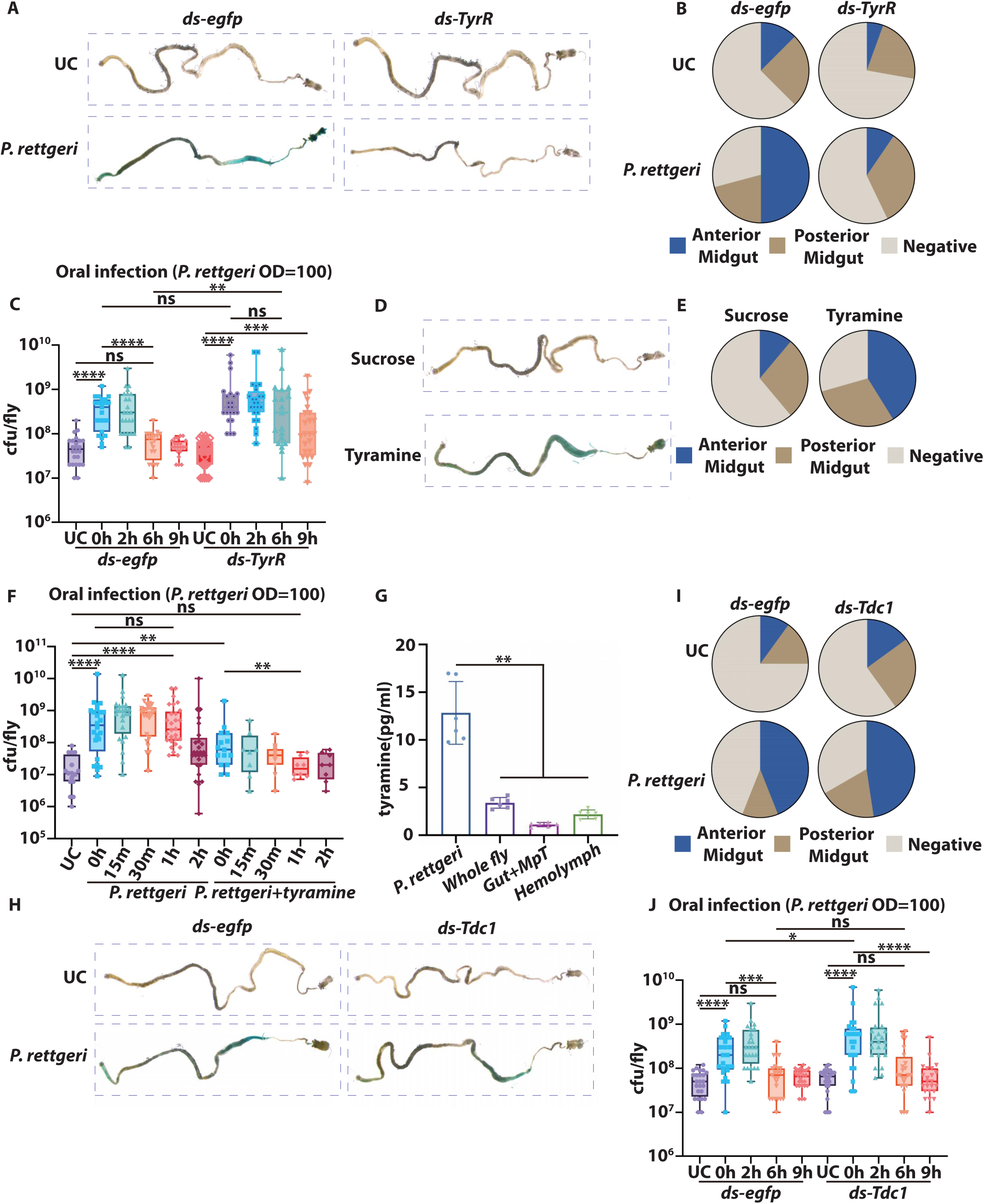
Exogenous tyramine is a signalling molecule that triggers MpT-gut countercurrent flow. **(A and B) *TyrR* RNAi reduces MpT-gut countercurrent.** *TyrR* RNAi flies showed very weak blue dye accumulation in the anterior midgut (A) and quantification in (B). *egfp* RNAi flies were used as control. (n=18-24). UC: unchallenged. **(C) *TyrR* RNAi decreased the flies’ ability to eliminate gut invading bacteria.** The Cfu counts in *TyrR* RNAi flies were significantly higher at 6 h compared to the control group. *egfp* RNAi flies were used as control. The data represent the average of 3 biological replicates (n=19-24). UC: unchallenged. **(D-F) Tyramine promotes MpT-gut countercurrent flow and bacteria clearance.** Tyramine-fed flies showed a strong accumulation of bright blue dye in the anterior midgut (D) and quantification in (E) (n=18 for sucrose, n=17 for tyramine). Co-fed tyramine and bacteria promoted the bacterial elimination (F) (n=8-24). The data represent the average of 3 biological replicates. UC: unchallenged. **(G) The concentration of tyramine in *P. rettgeri* culture and various tissues of *B. dorsalis*.** *P. rettgeri* produced higher amount of tyramine compared to that in the fly tissues. The data represent the average of 3 biological replicates (n=6). **(H-J) Endogenous tyramine did not affect MpT-gut countercurrent flow and bacteria elimination.** Silencing *Tdc1* did not affect the formation of MpT-gut countercurrent flow (H) and quantification in (I). *Tdc1* RNAi did not affect the clearance of *P. rettgeri* in the gut of *B.dorsalis* (J). *egfp* RNAi flies were used as control. The data represent the average of 3 biological replicates (n=20-25 for (H), n=23-24 for (J)). * p<0.05, ** p<0.01, *** p<0.001, **** p<0.0001, ns non-significant, Mann Whitney test. UC: unchallenged.

It is well characterized that bacteria, for example, *Providencia* bacteria, could produce large amounts of tyramine regulating host physiology^34^. We wondered if the tyramine was produced by the flies or if the bacteria secret it. We first examined the tyramine quantity in *P. rettgeri* and *B. dorsalis*. The results showed that both *P. rettgeri* and *B. dorsalis* could produce tyramine (Figure 3G). Notably, *P. rettgeri* produces much more tyramine than the amount in whole flies or various tissues, suggesting a possible role of bacteria-derived tyramine in regulating MpT functions. To validate our assumption, we knocked down the tyrosine decarboxylase 1 (*Tdc1*) in *B. dorsalis*. Tdc1 is the enzyme that catalyzes the decarboxylation of tyrosine to tyramine and is necessary for renal function regulation in *Drosophila*. The results showed that *Tdc1* RNAi flies had a similar MpT-gut countercurrent flow as *egfp* RNAi flies, and bright blue dye could reach the gut R2 region after *P. rettgeri* infection (Figures 3H and I), suggesting that intrinsic tyramine synthesis pathway is not required for countercurrent flow formation. Consistent with this finding, we also found that the bacteria clearance ability remained unchanged in *Tdc1* RNAi flies (Figure 3J). To further strengthen the conclusion, we detected tyramine levels in the gut and hemolymph in *Tdc1* RNAi flies after *P. rettgeri* infection. The results showed that *Tdc1* RNAi did not affect tyramine levels in the gut and hemolymph compared with the control group (Supplementary Figures 3A and B). In both tissues, we still observed an increase in tyramine content in both gut and hemolymph, suggesting that these tyramines are exogenous. Taken together, these data indicated that tyramine secreted by pathogenic bacteria activates the countercurrent flow.

### Duox-mediated ROS production, not antimicrobial peptides, ensures early-phase bacteria clearance in *Bactrocera*

Two inducible immune mechanisms, the production of ROS by Duox and AMP by the Imd pathway, have been involved in gut immunity in *Drosophila*. We first aimed to explore the mechanism involved in the clearance of bacteria in the early infection stage in *B. dorsalis*. We found that the level of *P. rettgeri* (Figure 4A, 0 h compared to UC) remained at a high level for the next hour upon return to fresh food. Strikingly, *P. rettgeri* bacterial count quickly returned to unchallenged suggesting that *B. dorsalis* cleared most invading bacteria (Figure 4A). Having demonstrated that *B. dorsalis* has potent gut immunity/mechanism to eliminate ingested bacteria, we next explore the role of AMPs and microbicidal ROS in this process.

**Figure 4.**
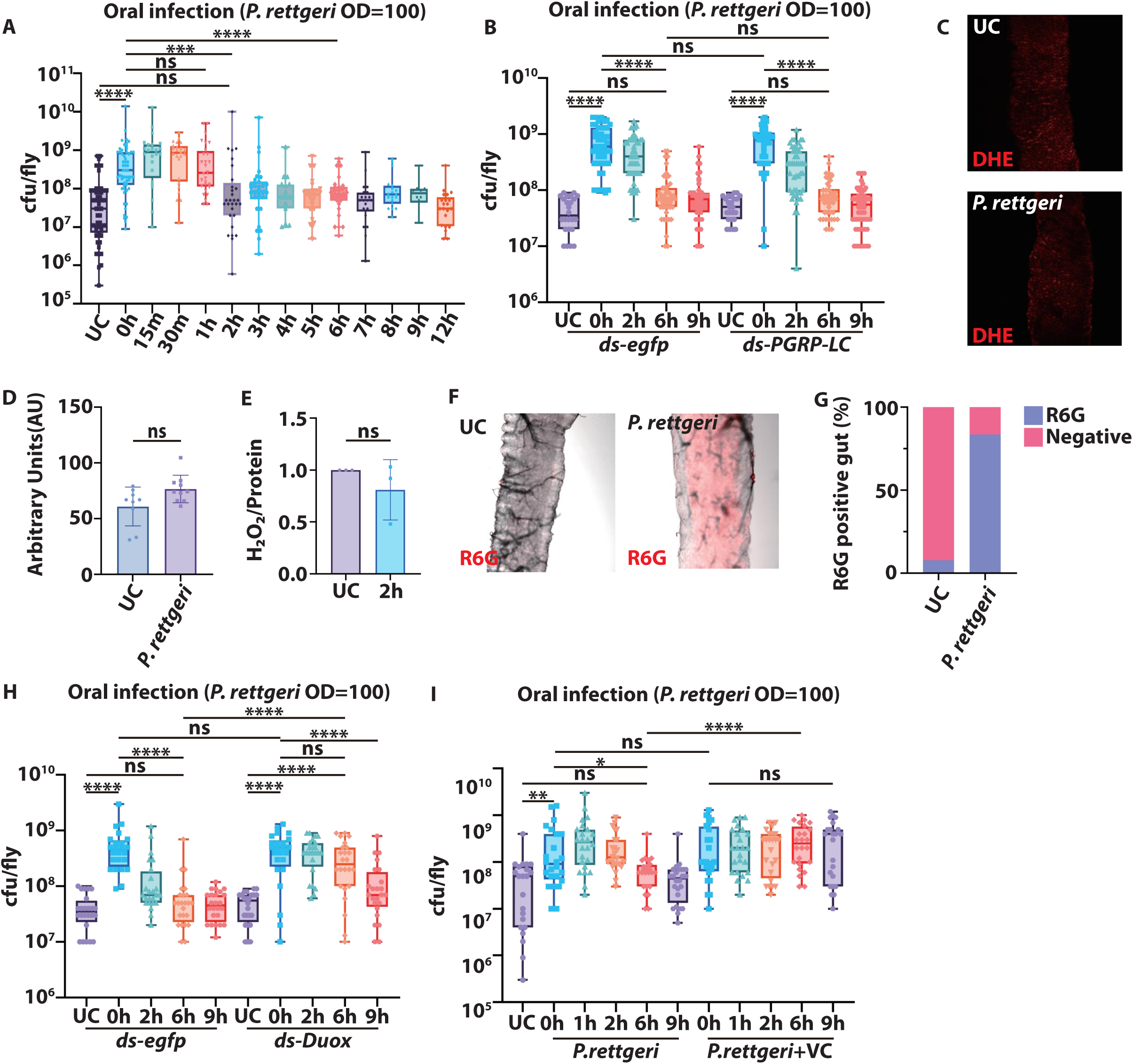
Duox-mediated ROS production but not the PGRP-LC IMD pathway ensures early-phase bacteria clearance. **(A) Bacteria persistence in *B. dorsalis* gut at different time points after gut infection.** The gut cfu counts were significantly lower at 2 h compared with 0 h after gut infection. The data represents the average of 3 biological replicates (n=12-56). UC: unchallenged. **(B) *PGRP-LC* RNAi did not affect bacterial elimination.** The gut cfu counts showed no difference in *PGRP-LC* RNAi flies compared to the control group. *egfp* RNAi flies were used as control. The data represents the average of 3 biological replicates (n=24-50). UC: unchallenged. **(C and D) Superoxide content in the gut after *P. rettgeri* gut infection.** Superoxide content in the gut was revealed by DHE staining (C) and quantification in (D). The data represents the average of 3 biological replicates (n=9 for UC and n=10 for *P. rettgeri*). UC: unchallenged. **(E) *P. rettgeri* gut infection did not activate gut H_2_O_2_ production.** The H_2_O_2_ content was normalized to the UC group, which was set as 1. The data represents the average of 3 biological replicates (n=3). UC: unchallenged. **(F and G) HOCl staining in the gut after *P. rettgeri* infection.** HOCl sensor R6G staining showed an increased amount of ClO^-^ in the gut after infection (F). The percentage of R6G-positive guts increased after infection (G). The data represents the average of 3 biological replicates (n=13 for UC, n=18 for *P. rettgeri*). UC: unchallenged. **(H and I) Removing gut ROS dampened *B. dorsalis* ability to eliminate invading bacteria.** Removing ROS production by *Duox* RNAi delayed bacteria elimination (H). Co-fed antioxidant VC with bacteria delayed *B. dorsalis* bacteria elimination (I). *egfp* RNAi flies were used as control. The data represents the average of 3 biological replicates (n=21-24). * p<0.05, ** p<0.01, *** p<0.001, **** p<0.0001, ns non-significant, Mann Whitney test. UC: unchallenged.

For this, we first checked the expression of AMPs after *P. rettgeri* infection. The results showed that *P. rettgeri* infection induced a strong up-regulation of *attacinA*, *attacinB*, *attcinC*, and *diptericin* genes, suggesting that the IMD pathway is activated upon gut bacterial infection (Supplementary Figures 4A-D). The Imd pathway is activated by the transmembrane receptor PGRP-LC, which recognizes peptidoglycan. Silencing PGRP-LC upon injection of dsRNA confirms that the IMD pathway was responsible for AMP expression in *B. dorsalis* (Supplementary Figures 4E-H). However, *PGRP-LC* RNAi flies showed normal bacteria clearance ability compared with *egfp* RNAi control flies (Figure 4B). This indicates that the Imd pathway does not play a critical role in bacteria elimination of bacteria.

We next explored the role of the Duox-ROS production system in bacterial clearance. We first checked ROS production in the gut after *P. rettgeri* infection 2 h post infection. The primary Duox ROS product is HOCl, while superoxide (O_2_^-^) and H_2_O_2_ are intermediate products. We monitored each of these ROS in the gut using appropriate kit assays: DHE for superoxide, hydrogen peroxide assay kit for H_2_O_2_ and the R6G hypochlorous acid sensor for HOCl. Interestingly, we did not detect any increase in superoxide production as indicated by DHE staining in the gut (Figure 4C and D), nor did we see an increased amount of H_2_O_2_ (Figure 4E). However, we saw a strong HOCl burst in the gut region corresponding to *Drosophila* region R2 at 2 h post-infection (Figure 4F and G)^35^. To check whether ROS is necessary for bacterial clearance, we monitored bacterial persistence in flies upon *Duox* RNAi. We found that *Duox* RNAi flies lost the ability to clear invading bacteria as these flies cannot remove the bacteria until 9 h after infection, suggesting that Duox-mediated ROS production mechanism is crucial for early *B. dorsalis* gut immune response (Figure 4H). To further support our conclusion, we co-fed the flies with bacteria and vitamin C, a potent antioxidant. We found that *B. dorsalis* cannot clear invading bacteria even at 9 h after infection when we remove the ROS using vitamin C (Figure 4I). Altogether, our data show that *Bactrocera* possess efficient mechanism to clear ingested bacteria iand that Duox-based ROS is the primary immune mediator involved early gut immunity in *B. dorsalis*.

### ROS production by Duox promotes bacterial clearance by peristalsis

Recent study in *Drosophila* has highlighted a possible key role of Duox in bacterial clearance by promoting muscle contraction^13, 14^. We therefore investigated whether peristalsis also contribute to *B. dorsalis* ability to eliminate bacteria. We fed flies with FITC-labelled dextran and monitored gut peristalsis upon oral infection with the Gram-negative bacteria *P. rettgeri* infection. We observed that oral infection increased gut peristalsis by 32% (Figure 5A). This effect was not observed in *Duox* RNAi flies indicated that in *Bactrocera* like in *Drosophila*, peristalsis is regulated by Duox (Figure 5B). Next, we monitored if peristalsis promote bacteria clearance. For this experiment, we used the rifampicin-resistant *Pectobacterium carotovorum* subsp. *carotovorum*-GFP (*Ecc15-GFP*) to track the bacteria in the faeces. We monitored the presence of live bacteria in the gut and faeces at 4 hours post-infection. We found six times more live bacteria in the faeces than in the gut after oral infection with *Ecc15-GFP*, indicating that in *B. dorsalis* too, gut peristalsis promotes bacteria purge (Figure 5C and 5D). Importantly, silencing *Duox* by RNAi suppressed the ability of *Bactrocera* to clear *Ecc15-GFP* by expulsion via the faeces (Figures 5E and 5F). Collectively, our study reveals that peristalsis requires Duox in *Bactrocera* and is critical for bacterial elimination.

**Figure 5.**
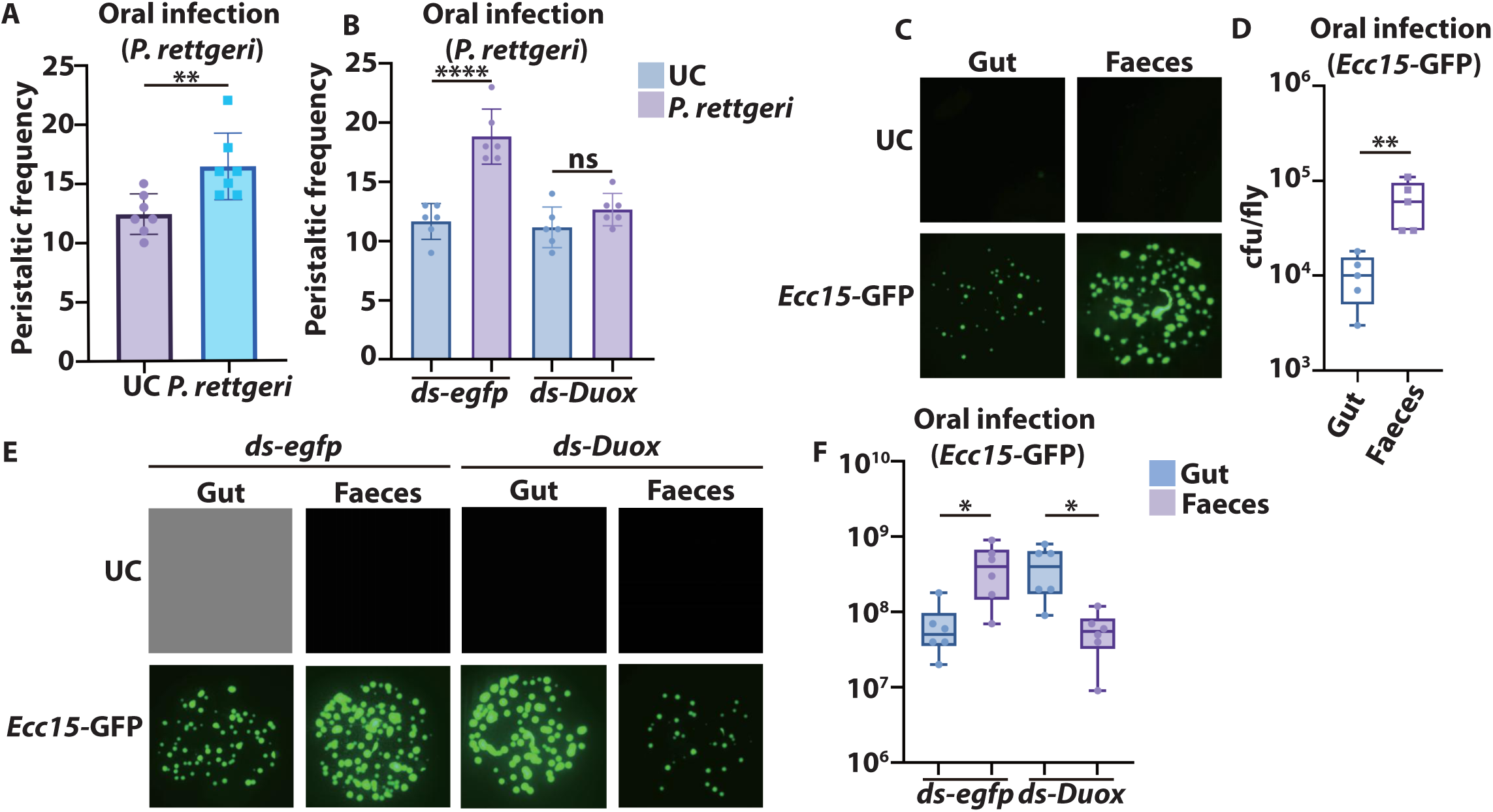
ROS production by Duox promotes bacterial clearance by peristalsis. **(A) The intestinal peristalsis frequency increased after *P. rettgeri* gut infection.** Two hours after *P. rettgeri* gut infection, the frequency of intestinal peristalsis increased significantly. The data represents the average of 3 biological replicates (n=7). UC: unchallenged. **(B) *Duox* RNAi decreases the intestinal peristalsis frequency after *P. rettgeri* gut infection.** Two hours after *P. rettgeri* gut infection, the frequency of intestinal peristalsis did not increase in *Duox* RNAi flies. The data represents the average of 3 biological replicates (n=6). UC: unchallenged. **(C) Invading bacteria are excreted after infection.** *Ecc15-*GFP was higher in the faeces than in the gut 2 h after infection. *Ecc15-*GFP is not observable in UC flies. UC: unchallenged. **(D) Cfu count of bacteria in the gut and faeces after infection.** The number of bacteria in faeces is higher compared with the number of bacteria in the gut. The data represents the average of 3 biological replicates (n=5). **(E-F) *Duox* RNAi delayed bacteria elimination and Cfu count of bacteria.** *Ecc15-*GFP was higher in the gut than in the faeces 2h after infection in *Duox* RNAi flies. *Ecc15-*GFP is not observable in UC flies. * p<0.05, ** p<0.01, **** p<0.0001, Mann Whitney test. UC: unchallenged.

### ROS are maintained in the midgut by a countercurrent flow

We expected that gut peristalsis would also induce a strong clearance of gut ROS, which should inhibit peristalsis-mediated bacteria clearance. We, therefore, checked the H_2_O_2_ amount, one of the intermediate reactive oxygen products of Duox, in the gut and the faeces. Although we saw only a mild increase in fecal ROS amount after bacteria oral infection, most of the ROS were found in the gut (Figure 6A). Thus, *B. dorsalis* has a mechanism to preserve the ROS within the gut despite peristalsis.

**Figure 6.**
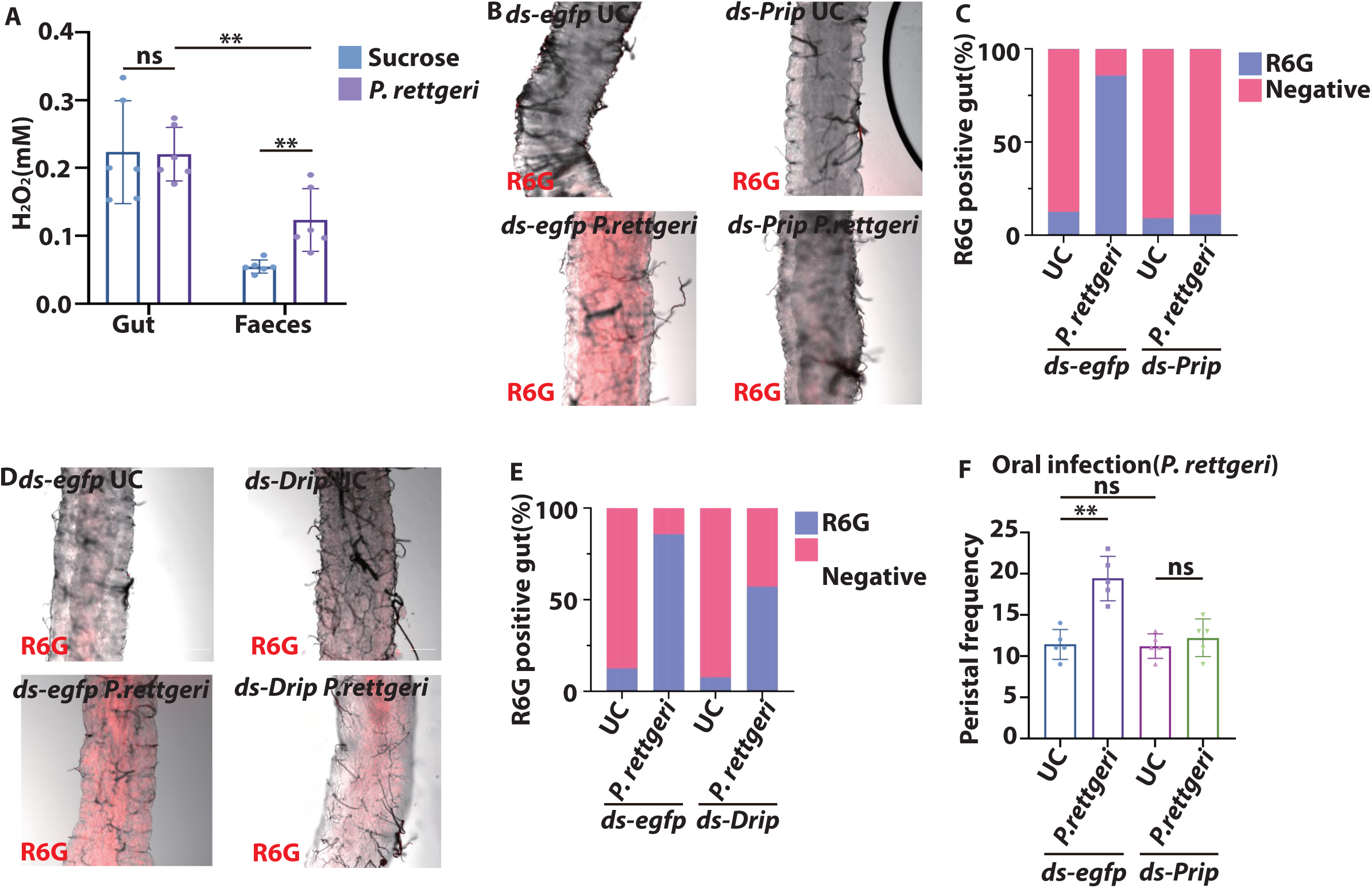
Countercurrent promotes ROS accumulation and gut peristalsis. **(A) H_2_O_2_ concentration in the gut and faeces of *B. dorsalis* after *Ecc15-*GFP infection.** The H_2_O_2_ was lower in the faeces compared to the gut. The data represents the average of 3 biological replicates (n=6). ** p<0.0, ns non-significant, Mann Whitney test. UC: unchallenged. **(B and C) *Prip* RNAi reduced the ClO^-^ accumulation in the gut.** Only *egfp* RNAi flies showed a strong gut R6G signal after *P. rettgeri* oral infection (B) and quantification in (C). *egfp* RNAi flies were used as control. (n=9-16). UC: unchallenged. **(D and E) *Drip* RNAi did not affect the ClO^-^ accumulation in the gut.** Both *egfp* RNAi and *Drip* RNAi flies showed a strong R6G fluorescence signal in the gut after P. *rettgeri* oral infection (D) and quantification in (E). *egfp* RNAi flies were used as control. (n=13-16). UC: unchallenged. **(F) *Prip* RNAi reduces the gut peristalsis frequency.** *Prip* RNAi flies did not increase gut peristalisis after infection. *egfp* RNAi flies were used as control. The data represent the average of 3 biological replicates (n=5). ** p<0.01, ns non-significant, Mann Whitney test. UC: unchallenged.

We wondered whether blocking MpT-gut countercurrent flow would affect the ROS amount in the gut. We detected the HOCl amount using *Prip* RNAi flies. The results showed that *Prip* RNAi strongly decreased the HOCl amount in the gut lumen after oral infection, as indicated by the R6G HOCl sensor (Figure 6B). Quantification showed that the R6G-positive gut number in the infected group was similar to the unchallenged group in *Prip* RNAi flies (Figure 6C). This result demonstrated that countercurrent flow promotes gut bacteria clearance through HOCl accumulation. To further support our conclusion, we also detected HOCl amount in *Drip* RNAi flies. As expected, *Drip* RNAi did not affect ROS amount in the gut, suggesting that MpT-gut countercurrent flow is essential for preventing ROS purge (Figures 6D and 6E). To exclude the possibility that *Prip* RNAi would directly affect AMPs and ROS production. We examined *Diptericin* and *Duox* expression in *Prip* RNAi flies. The qRT-PCR analysis showed that *Prip* RNAi did not affect the *Diptericin* and *Duox* expression (Supplementary Figures 5A and 5B). Since Duox played an important role in gut muscle contraction and food defecation, we checked gut peristalsis in *Prip* RNAi flies. Interestingly, *Prip* RNAi inhibited the gut peristalsis increase after infection (Figure 6F). Taken together, these observations suggested that the MpT-gut countercurrent flow ensures gut pathogenic bacteria clearance by promoting ROS-mediated gut peristalsis.

### MpT-gut countercurrent flow is necessary for microbiota regulation

Many bacteria besides pathogenic bacteria could produce tyramine. We wondered whether gut microbiota could produce tyramine and induce countercurrent flow as *P. rettgeri*. We examined tyramine quantity in four major *B. dorsalis* microbiota species cultures^30^. We found that they also produced a comparable amount of tyramine to *P. rettgeri* (Figure 7A, Compared with Figure 3G). Based on our theory, tyramine production by microbiota could have a critical effect in regulating microbiota composition. We performed 16S rRNA sequencing in *Prip* RNAi flies to validate this assumption. Regarding relative abundance, the high-throughput sequencing revealed that *Enterobacteriaceae*, *Morganellaceae*, *Orbaceae*, and *Streptococcaceae* dominated the gut regions of *egfp* RNAi control flies. However, *Enterobacteriaceae*, the most abundant bacteria species, decreased to 22.48% in *Prip* RNAi flies compared to the control group. The less dominant species *Streptococcaceae* gradually reduced to very low levels, becoming a minor microbiota species in the flies’ gut. On the contrary, we found a substantial increase in *Morganellaceae*, which accounts for around 35.39% of the total microbiota in *Prip* RNAi flies. One minor species in control flies, *Desuifovobrionaceae*, also became a dominant microbiota species in *Prip* RNAi flies (Figure 7B). These results suggested that inhibiting MpT-gut countercurrent flow by *Prip* knockdown caused microbiota dysplasia in the gut. Furthermore, gut microbiota species richness and evenness were reduced in the *Prip* RNAi flies, as indicated by Simpson diversity indices (Figure 7C).

**Figure 7.**
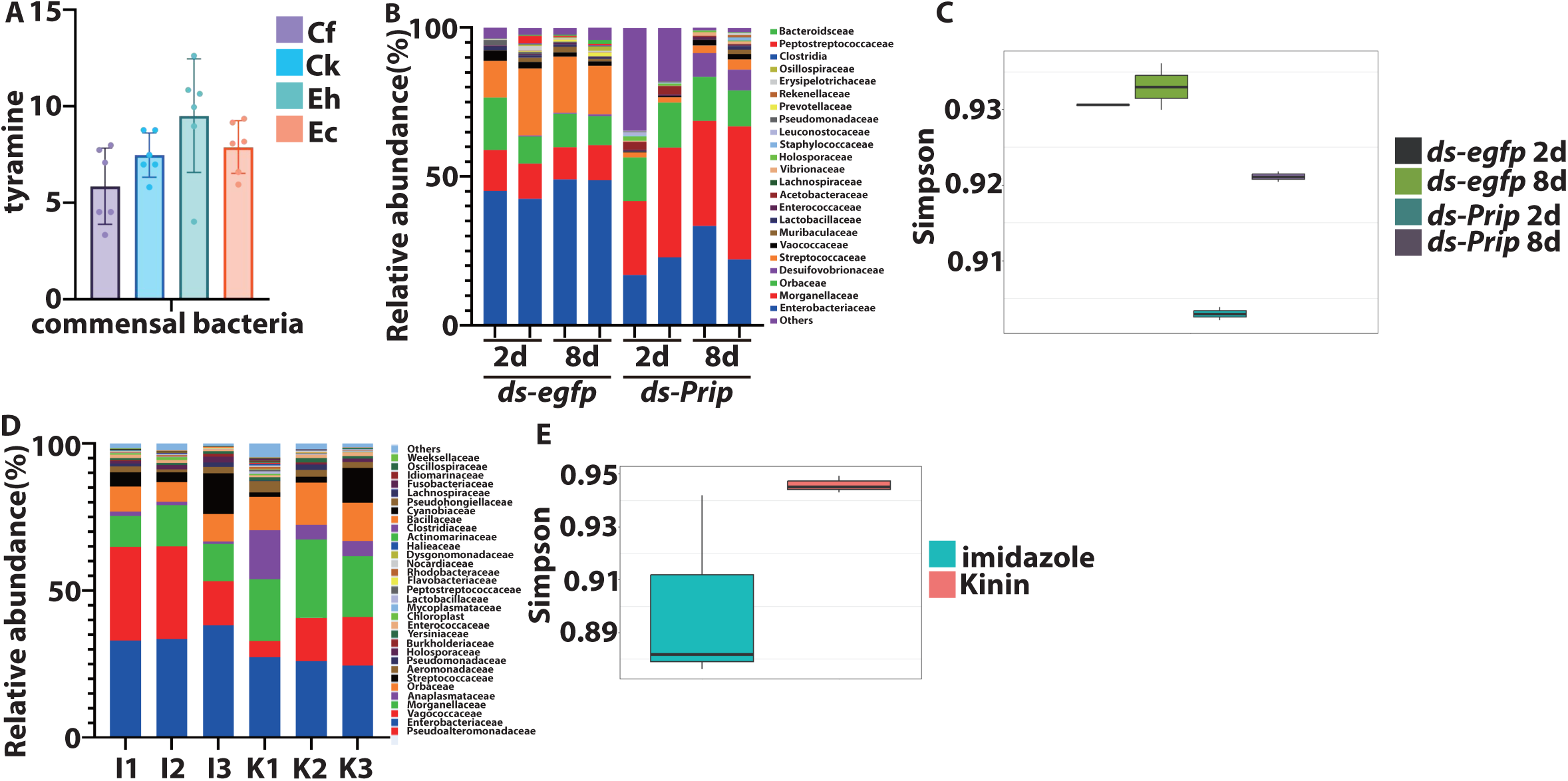
MpT-gut countercurrent flow is necessary for microbiota regulation. **(A) The commensal bacteria also produced tyramine.** Cf: *Citrobacter freundii*, Ck: *Citrobacter koseri*, Eh: *Enterobacter hormaechei*, Ec: *Enterobacter cloacae*, The data represent the average of 3 biological replicates (n=6). **(B and C) *Prip* RNAi altered gut microbiota composition in *B. doralis*.** Relative genus-level abundance profiles of bacteria in the gut (B). Gut microbiota community diversity was measured using the Simpson index (C). *egfp* RNAi flies were used as control. (n=2). **(D and E) Kinin injection altered gut microbiota composition in *B. doralis.*** Relative genus-level abundance profiles of bacteria in the gut (D). Gut microbiota community diversity was measured using the Simpson index (E). (n=3). UC: imidazole.

To further strengthen the conclusion, we performed 16S rRNA sequencing of flies injected with kinin. Regarding relative abundance, the high-throughput sequencing revealed that *Enterobacteriaceae*, *Vagococcaceae*, *Morganellaceae*, *and Orbaceae* dominated the gut regions of control flies. However, *Enterobacteriaceae*, the most abundant bacteria species, decreased to 8.9% in kinin injection flies compared to the control group. The less dominant species *Vagococcaceae* gradually reduced to lower levels, becoming a minor microbiota species in the flies’ gut. On the contrary, we found a substantial increase in *Morganellaceae*, which accounts for around 10.4% of the total microbiota in kinin injection flies. One minor species in control flies, *Anaplasmataceae*, also became a dominant microbiota species in kinin injection flies (Figure 7D). According to Simpson diversity indexes, the flies injected with kinin had higher bacteria diversity (Figure 7E). Altogether, these data supported the notion that MpT-gut countercurrent flow regulates gut microbiota composition and maintains its stability.

## Discussion

In this study, we identified a physiological mechanism essential for gut immune response and microbiota homeostasis. We demonstrated that Duox-derived ROS-promoting gut peristalsis is the major driving force for pathogenic bacteria elimination. The host generates a MpT-gut countercurrent flow in response to the presence of bacteria-produced tyramine. This flow counteracts the forced defecation of the gut peristalsis and drives the accumulation of ROS. We also provided evidence that this flow is required to maintain microbiota homeostasis. This work demonstrated a mechanism promoting the recycling or accumulation of antimicrobial agents in insects despite the increased gut peristalsis. Furthermore, our work also proved the importance of MpT in the gut defense mechanisms, in addition to their well-known functions in regulating excretion physiology.

The fact that ROS is well preserved by the MpT-gut countercurrent flow proved that insects own a delicate system to prevent the loss of valuable resources. The countercurrent flow from MpT to the anterior gut in *B. dorsalis* appears to be conserved in other insects. Recently, we found that countercurrent flow induced by *Ecc15* infection was crucial for gut epithelial renewal in *D. melanogaster*^26^. Early reports in locusts showed that a counter-current flow from MpT occurred when the locusts were starved^28^. By using dye feeding and observation of its distribution in the gut, Terra and colleagues proposed the existence of countercurrent flow in other insects, including Lepidoptera ^36^, Coleoptera^37^, and Orthoptera^38^. Our results suggested aquaporins’ critical role in forming the countercurrent flow in *B. dorsalis*. Consistent with this, we found that *Drip* in stellate cells was required for MpT-gut retrograde flow formation in *Drosophila*^26^. Interestingly, aquaporins are also enriched in the anterior midgut in *Drosophila*, implying that the anterior midgut might be a major region for water absorption, thus forming a complete water flow cycle consisting of MpT, hemolymph, and the gut^35, 39^.

The originally proposed function of countercurrent flow is the recycling of enzymes to increase digestion efficiency. The most solid evidence is that the enzyme concentration is higher within the ectoperitrophic space than in the endoperitrophic space^29^. Our previous results suggested that this countercurrent flow could bring renal-derived Upd3 to regulate gut renewal upon oral infection^26^. Here, we found that the retroflow also participated in the immune response by promoting ROS accumulation and gut peristalsis. It is reasonable to expect that this mechanism could also promote the accumulation of other antimicrobial agents, such as AMPs, or regulate the immune-metabolism relationship.

Our results suggested that gut peristalsis is crucial for pathogenic bacteria and microbiota control. The increased gut motility in response to pathogens is a conserved response. Animal hosts actively sequester bacteria or parasites within the intestinal lumen to protect mucosal tissues and maintain immune tolerance. In mammals, parasitic nematode *Trichinella spiralis* increases intestinal motility and propulsion as the primary mode of elimination, which is achieved by IL-33-induced instantaneous peristaltic movement ^40–42^. It has been shown that gut microbial products regulate murine gastrointestinal motility via Toll-Like receptor 4 signaling^1^. Other reports indicated that TLR2 is also possibly involved in this process^43, 44^. In *Drosophila*, this process involved TRPA1 sensing HOCl in a Duox-dependent manner and release of Dh31 in enteroendocrine cells. Our results also confirmed the critical role of gut peristalsis in gut immunity. We found that blocking gut peristalsis by knocking down *Prip* decreased flies’ bacteria clearance ability. We observed a considerable number of bacteria within the gut at 9 h post-infection in *Prip* RNAi flies, suggesting that gut peristalsis is crucial for pathogen elimination. Similar to our experiment, Du et al. reported that opportunistic bacteria *Ecc15* induce accelerated defecation between 2 h and 8 h post 2 h infection in *D. melanogaster*^13^. Interestingly, a short opportunistic pathogens infection (30 mins) induces a shorter gut contraction (less than 4 hours)^14^. Therefore, we can assume that continuous bacterial infection will cause prolonged gut peristalsis until the pathogen is eliminated. Altogether, these data suggested that increased gut peristalsis is the primary driving force toward accelerated and successful bacteria elimination.

ROS generation is highly conserved across all organisms as a physiological response to microbes^45^. Duox-derived ROS has been proposed to have a direct bacteria-killing effect in gut immunity and microbiota maintenance^20, 21, 23^. *In vitro* biochemical evidence supports the role of Duox-derived HOCl but not H_2_O_2_ as the major microbicidal molecule in *Drosophila* gut immunity^20^. This direct microbicidal role of Duox in disease resistance was also implied by studies in mosquitos and other insects^46, 47^. However, Duox has also been suggested to be indirectly involved in gut immunity, possibly as a signaling molecule. For example, Duox-dependent ROS is involved in epithelial cell renewal during gut infection^25, 26, 48^ and indispensable for gut muscle contraction and food defecation in *Drosophila* during gut infection^13, 14^. Recently, it was shown that Duox is also indirectly involved in regulating gut symbionts in the bean bug *Riptortus pedestris*^49^. In this bug, *Duox* expression is tracheal specific and necessary for tracheal integrity, which is crucial to sustain mutualistic symbionts. These data together suggested a dedicate and controversial role of Duox in insect gut immunity. Notably, in neutrophil phagosomes, HOCl is expected to have a lifetime of 0.1 μs and a diffusion length of ∼30 nm^50, 51^. However, the maximum peritrophic membrane thickness was variable from 2µm to 20µm depending on insects, not to mention the distance of ectoperitrophic space^5, 52^. This short lifetime and reach would confine HOCl working distance. Furthermore, the previous conclusion was drawn from a continuous bacterial oral infection, which might be a mixed outcome of both disease resistance and tolerance^20^. Our experiments were done using a short-time oral infection and only focused on the bacteria elimination in the early stage of infection. We found that ROS depletion inhibits bacteria elimination due to Duox’s role in regulating gut peristalsis, similar to previous findings^13, 14^. More importantly, we found that *Duox* RNAi only slowed down the clearance of bacteria but did not completely inhibit the process. Unlike our observation, Ha et al. reported a prolonged *Ecc15-GFP* persistence in the *Duox* RNAi *Drosophila* gut at 60 h post-infection, which could be explained by continuous bacteria feeding and long-lasting inhibition of gut peristalsis, as we discussed earlier. Thus, we propose that early induction of ROS played a major role in supporting gut peristalsis to accelerate food-borne bacteria elimination. However, we cannot exclude the possibility that ROS accumulated in the gut plays a direct bactericidal function, especially under continuous infection. Whether HOCl has a direct microbicidal effect on insect gut immunity remains to be further investigated.

In conclusion, our study has clarified the interaction of Duox and gut peristalsis in gut immune response. Importantly, we have characterized MpT as a key organ regulating gut immunity through recognizing bacteria-derived tyramine, providing an example of inter-organ communication in gut immunity. Recently, two papers have shown that gut tumors and MpT have complex interactions promoting tumor-induced renal dysfunction^53, 54^. Since MpT plays a key role in detoxification and excretion, we expect more of the role of the renal system in host physiology will be revealed in the future.

## Acknowledgment

This study was supported by National Key R&D Program of China (No. 2021YFC2600400), China Agriculture Research System of MOF and MARA (CARS-26) and Hubei Hongshan Laboratory. We thank Weiwei Zheng for her insightful comment on the manuscript.

## Author contributions

X.L. conceived and designed the experiments, Y.L. and R.L performed the experiments, Y.L. and X.L. analyzed the data, X.L., B.L., H.Z. and Y.L. wrote the manuscript, X.L. and H.Z. supervised the project.

## Materials availability

Further information and requests for resources and reagents should be directed to and will be fulfilled by the lead contact, Xiaoxue Li.

## Methods

### Insect rearing

The oriental fruit flies were reared at the Institute of Horticultural and Urban Entomology, Huazhong Agricultural University (Wuhan, China), with a 14 h light/10 h dark cycle at 27 ± 1 ℃ and 70–80 % relative humidity. Larvae were raised on an artificial diet constituting of 300 g banana, 300 g corn meal, 60 g yeast, 60 g sucrose, 1.2 g sodium paraben methyl ester, 60 g tissue paper, 2.4 ml hydrochloric acid, 600 ml water. After eclosion, adult flies were moved to 30 × 30 × 30 cm cages and maintained on an artificial diet. The artificial diet made of 10 % sucrose, 3.4 % yeast, 1 % agar, 1.68 % honey, supplemented with 1.6 g/L sodium paraben methyl ester. Female flies that emergence for 5 to 8 days were used for all experiments. *Drosophila* stocks were maintained at 25 °C on standard fly medium made of 6 % cornmeal, 6 % yeast, 0.62 % agar, 0.1 % fruit juice, supplemented with 10.6 g/L moldex and 4.9 ml/L propionic acid.

### Oral infection and Bacteria persistence

*Providencia rettgeri* were cultured in Luria-Bertani (LB) medium overnight (14 h) at 37 °C 220 rpm from single colony. *Pectobacterium carotovorum* subsp. *carotovorum* (*Ecc15*-GFP) were cultured in Luria-Bertani (LB) medium overnight (14 h) at 29 °C 220 rpm from single colony. Bacteria culture were harvested by centrifuge (4200 rpm, 4 °C, 10 min). The infection solution was obtained by mixing an equal volume of an overnight culture of *P. rettgeri* (optical density OD600 = 200) with a 5 % sucrose solution (1:1) covering the surface of the artificial diet. Five to eight days old adult female *B. dorsalis* were starved in an empty box at 28 °C for 2 h, then supplied with the infection solution. For all experiments in this paper, the flies were subjected to a 2h infection and then switched to a clean fly food for the following sampling. The time points mentioned in the paper indicate the time post the 2h infection process. For the bacteria persistence experiment, the flies were firstly washed twice in 75 % ethanol for 30 s, then washed using PBS for 30 s. The flies were then homogenized in a 2 ml freezing tube containing 200 μl of LB solution and 3 steel beads. For faeces samples collection, flies were moved to a 50 mL falcon tube after infection and then faeces were washed and collected using LB at desired time point. The samples were serially diluted and dilutions were then plated (2 μl) onto the LB plates. The plates were cultivated overnight at 37 °C (*P. rettgeri*) and 29 °C (*Ecc15*-GFP). *Ecc15*-GFP plates were imaged using stereo-fluorescence microscope MZX81 (MshOt, China). For vitamin C (VC) feeding experiment, 50 mg/ml of VC was added to the infection solution. For tyramine feeding experiment, 70 mM of tyramine was added to the infection solution.

### Monitor of intestinal peristalsis frequency

Five to eight days old adult female *B. dorsalis* were starved in an empty box at 28 °C for 2 h. The flies were then infected with infection solution supplied with FITC dextran. One minute gut peristalsis videos were recorded using stereo-fluorescence microscope MZX81 (MshOt, China) 2 h after infection. The gut peristalsis times were counted from the videos.

### Gut ROS staining and quantification

For ROS measurement with DHE, flies were anesthetized on the ice and gut were dissected in Schneider’s insect medium. The guts were directly incubated in 30 mM DHE (Life Technologies) for 7 min at room temperature, washed twice with Schneider’s insect medium and imaged immediately by a SP8 LIGHTNING confocal microscope. The signal intensity was quantified using FIJI.

R6G hypochlorous acid sensor was purchased from Heliosense (Xiamen, China). For microscope scanning of the R6G fluorescence signal, 50 μM of R6G was added to the infection solution and sucrose control solution. Flies were allowed to feed for 2 h on the solution. Flies were then collected 2h after the infection process and dissected in the Schneider’s insect medium. Guts were mounted in an antifade mounting medium (Beyotime, China) and immediately observed on a SP8 LIGHTNING confocal microscope.

For detecting H_2_O_2_ in the gut, we used hydrogen peroxide assay kit following the manufactural protocol. Briefly, flies were anesthetized on the ice and the guts were dissected in PBS at desired timepoints. The kit is based on the production of trivalent iron ions by oxidizing divalent iron ions with hydrogen peroxide, which then forms a purple product with xylenol orange. The absorbance was measured at 560 nm. H_2_O_2_ concentration was adjusted to the protein quantity using BCA assay.

### RNA interference

Specific dsRNA primers with the T7 RNA polymerase promoter (5’-GGATCCTAATACGACTCACTATAGG-3’) on the 5’ end were used to clone the target sequence fragments by PCR. Sequences of the primers were listed in Table S3. One μg PCR product was used as the specific-template for synthesizing of dsRNA in vitro by using the T7 Ribomax Express RNAi System (Promega, Madison, WI, USA). The concentration of dsRNA was quantified at 280 nm using a NanoDrop 2000 Spectrophotometer (Thermo Fisher Scientific Inc.). The quality and integrity of dsRNA were determined by agarose gel electrophoresis. Needles for injecting dsRNA were produced by a puller (PC-10, Narishige, Tokyo, Japan) at heat level 60.8. Microinjection of dsRNA was performed by Nanoinject III (Drummond Scientific, USA). RNAi experiments were performed by injecting 1 μL of a 2 μg μL^-1^ dsRNA into the fly thorax (5–8 days old). Experiments were performed 2-5 d after injection. Ds-egfp injected flies was used as control group.

### RNA extraction and quantitative PCR

For gene expression analysis, 8 whole flies or 20 guts were collected for RNA extraction. 500 ng total RNA or its tissues were then reverse-transcribed using PrimeScript RT (TAKARA) and a mixture of oligo-dT and random hexamer primers. Quantitative PCR was performed on cDNA samples in 20 μL reaction volume included 10 μL of SYBR Green Mix (Vazyme, China), 400 nM of each primer and 2 μL of cDNA (diluted 1:10) on an FQD-96X (Bioer, China) in 96-well plates with the following protocol: initial denaturation of 95 °C for 3 min, followed by 40 cycles of 95 °C for 15 s and 60 °C for 30 s. Relative quantification was calculated according to the 2^-ΔΔCt^ method. At least three independent biological replicates were performed. RpL32 was used as reference gene. Primers used in qPCR analysis were listed in Table S3.

### Countercurrent flow measurement

Two small wells were dug next to each other in a polydimethylsiloxane (PDMS) plate. A 1:1 mixture of Schneider’s insect medium and insect saline buffer was added into the two wells. Brilliant Blue solution was added into one of the wells to make the final concentration of 0.5 g/L. The gut and Malpighian tubules (MpT) of *B. dorsalis* were dissected. The MpT was put into the well with Brilliant Blue dye, and the gut was put into the other well. The PDMS plate was then covered with mineral oil to protect the tissue from evaporation. The gut was imaged under the microscope after 2 h.

### Leucokinin expression and injection

The whole Leucokinin ORF sequence was cloned into pET-N-His-TEV. E. coli HT115 (DE3) strain was transformed using pET-N-His-TEV-Leucokinin. The bacteria were incubated overnight at 37 °C. The overnight culture was diluted at 1:100 using LB and cultured until OD600 = 0.6. Then isopropyl β-D-thiogalactoside (IPTG) was added at final concentration of 0.1 mM, followed by overnight incubation at 18 °C 100 rpm. Mechanical lysis by sonication was performed on ice using sonic dismembrator with 200 hz for 30min. Bacteria lysis was purified His-tag protein purification kit (Beyotime, China) according to the manufactural protocol. The purified protein was verified by SDS-PAGE. Microinjection of Leucokinin was performed by Nanoinject III (Drummond Scientific, USA). 1 μL of Leucokinin was injected into the fly thorax (5– 8 days old). Experiments were performed 2-5 d after injection. Imidazole injected flies was used as control group.

### Hemolymph extraction and tyramine detection

For collecting hemolymph, 8 individuals were placed in a bottom-opened 0.6 ml tube covered with glass beads. This tube was put into a 1.5ml tube and centrifuged for 10 min at 4 °C, 1500 rpm. Debris were removed by centrifugation for 5 min at 13000 rpm for hemolymph samples. For gut and MpT tyramine detection, the desired tissues were dissected in PBS. The whole flies, gut, MpT and hemolymph samples were homogenized in PBS. For bacteria tyramine detection, overnight culture (OD600 = 100) was used. Tyramine detection was performed using insect tyramine kit (MEMIAN, China) according to the manufactural protocol.

### Bacteria DNA extraction and 16S rRNA amplicon sequencing

Bacteria DNA extraction and 16S rRNA amplicon sequencing were performed using the established method in the lab. DNA was extracted from 40 gut of flies (age: 5-8 days after emergence) per biological replicate using the E.Z.N.A. Soil DNA kit (Omega, Norcross, GA, USA) according to the manufacturer’s instructions, two or three biological replicates were conducted. The 16S rRNA gene spanning variable regions V3+V4 was amplified using the broad-range forward primer 341F: CCTAYGGGRBGCASCAG and the reverse primer 806R: GGACTACNNGGGTATCTAAT using the Phusion® High-Fidelity PCR Master Mix (New England Biolabs, Beverley, MA). The PCR amplification program consisted of (1) preincubation at 95 °C for 5 min; (2) 35 cycles of 45 s at 56 °C, then 1 min at 72 °C and 45 s at 94 °C; (3) 10 min at 72 °C. The PCR products were sent to Novogene Experimental Department for sequencing. After obtaining the raw reads, the single-end reads were assigned to samples based on their unique barcode and truncated by cutting off the barcode and primer sequence. Quality filtering on the raw reads was performed under specific filtering conditions to obtain high-quality clean reads according to the Cutadapt (b1.9.1, http://cutadapt.readthedocs.io/en/stable/) quality-controlled process (Martin, 2011). The reads were compared with the reference database (Silva database, https://www.arb-silva.de/) using UCHIME algorithm (UCHIME Algorithm, http://www.drive5.com/usearch/manual/uchime_algo.html) to detect chimera sequences, and then the chimera sequences were removed, the Clean Reads finally obtained. Alpha diversity (simpson) and beta diversity on weighted unifrac were calculated with qime and displayed with R software.

### Statistical analysis

Each experiment was repeated independently a minimum of three times (unless otherwise indicated). Error bars represent the standard deviation of the mean of replicate experiments. Data were analyzed using appropriate statistical tests as indicated in the figure legends using the GraphPad Prism software. p-values are represented in the figures by the following symbols: ns for P≥0.05, * for P between 0.01 and 0.05; ** for P between 0.001 and 0.01, *** for P between 0.0001 and 0.001, **** for P≤0.0001.

