## Supplementary Figures for "Bacterial-tyramine-induced renal-gut countercurrent flow regulates gut immune response by promoting ROS accumulation and gut peristalsis in the fly *Bactrocera dorsalis*"

**Supplementary Figure 1**

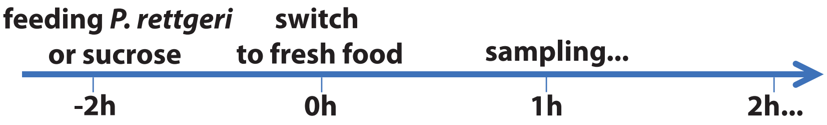

### Supplementary Figure 2

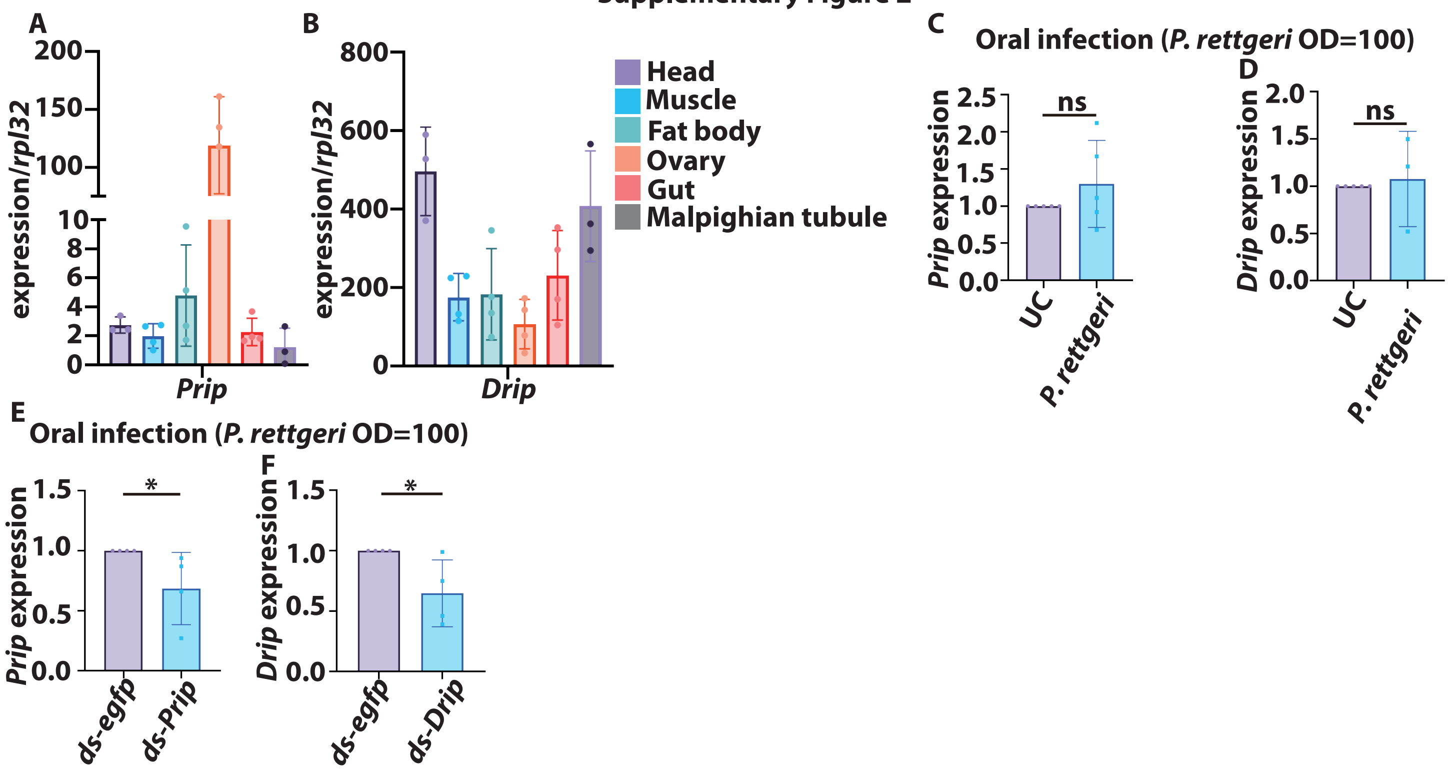

Supplementary Figure 3

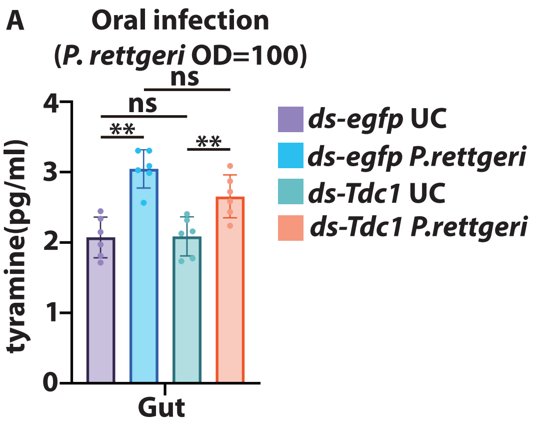


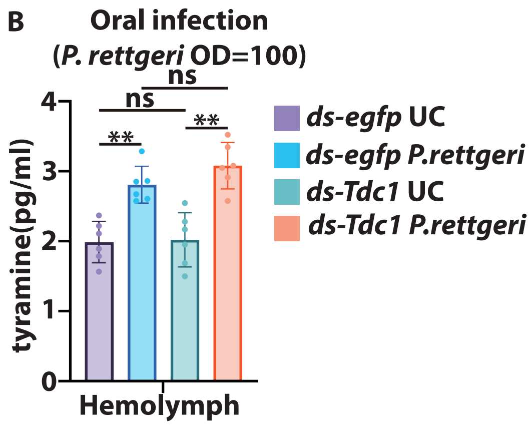

### Supplementary Figure 4

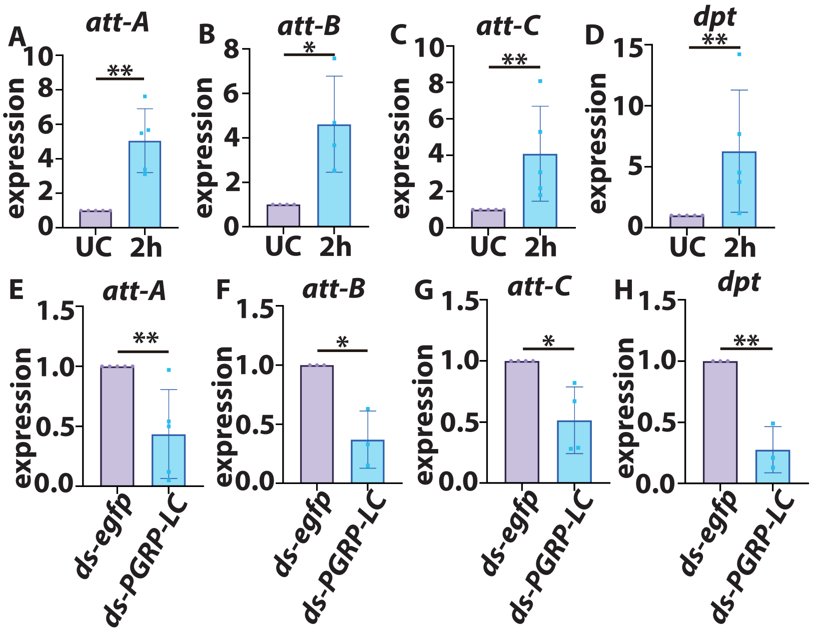

**Supplementary Figure 5**

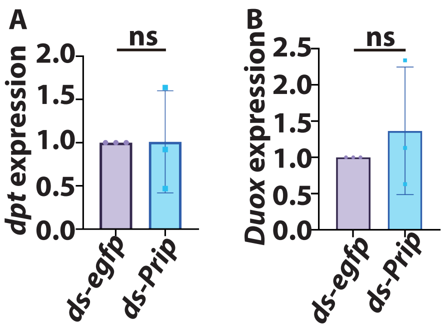
